# Masculinizer-induced dosage compensation is achieved by transcriptional downregulation of both copies of Z-linked genes in the silkworm, *Bombyx mori*

**DOI:** 10.1101/2022.03.06.483149

**Authors:** Kenta Tomihara, Munetaka Kawamoto, Yutaka Suzuki, Susumu Katsuma, Takashi Kiuchi

## Abstract

Dosage compensation balances the expression of sex-chromosome-linked genes with autosome-linked genes in the heterogametic sex. The silkworm (*Bombyx mori*), a lepidopteran model insect, uses the female heterogametic WZ sex determination system. In *B. mori*, a Z-linked gene, *Masculinizer* (*Masc*), is the primary determinant of maleness and dosage compensation. However, it remains unknown whether one of the two Z chromosomes is inactivated or both Z chromosomes are suppressed in *B. mori* males. Hence, we performed transcriptome analysis using hybrids between two *B. mori* strains and analyzed the allele-specific expression (ASE) to solve the problem. The ASE analysis revealed that the genes located on the maternal and paternal Z chromosomes are transcriptionally upregulated in *Masc* knocked down males. Also, our results revealed that two Z chromosomes are transcriptionally downregulated in *B. mori* males as observed in *Caenorhabditis elegans*.

## Introduction

Dosage compensation normalizes sex chromosome-linked gene expression with autosomes. Although dosage compensation is observed in many species, the mechanism has highly diverged. Therian mammals, the nematode *Caenorhabditis elegans* and the fruit fly *Drosophila melanogaster* use male heterogametic (XY or XO) sex determination systems. Therian mammals widely use the XY determination system. Dosage compensation is achieved by randomly silencing one of the two X chromosomes in female cells in eutherians, a lineage of therian mammals, including humans. [1]. For example, females heterozygous for X-linked coat colour mutations (e.g., *mottled*) in the mouse *Mus musculus* exhibit mosaic coat patterns [2]. In metatherians, another lineage of therian mammals, paternally inherited X chromosome is inactivated in females relying on genomic imprinting [3]. However, genes on the X chromosome are hyper-transcribed in males to match the expression in XX females in *D. melanogaster*, which also uses XY sex determination system [4,5]. Also, in *C. elegans*, which uses the XO sex determination system, the expression of X-linked genes in XX hermaphrodites are suppressed to match the expression from the single X in males [6].

The silkworm (*Bombyx mori*), a model species of lepidopteran insect, uses a female heterogametic (WZ) sex determination system [7]. Kiuchi et al. (2014) reported that a Z-linked gene, *Masculinizer* (*Masc*), is the primary determinant of maleness and dosage compensation [8]. Most Z-chromosome-derived transcripts are differentially expressed in male embryos injected with *Masc* siRNA [8]. This demonstrates that the Masc protein globally represses gene expression from the male Z chromosome at the embryonic stage. A similar Z-linked dosage compensation is widely observed in other lepidopteran species, such as *Cydia pomonella, Manduca sexta, Heliconius melpomene, H. cydno, Papilio xuthus, P. machaon* and *Plodia interpunctella*, with various developmental stages and tissues [9–12]. However, it remains unknown whether randomly silencing one of the two Z chromosomes (“eutherian-like” dosage compensation), silencing one of the two Z chromosomes depending on their parental origin (“metatherian-like” dosage compensation), or transcriptional downregulation of both Z chromosome copies (“nematode-like” dosage compensation) effectively achieved dosage compensation in lepidopteran males. Recently, Rosin et al. (2021) visualized *B. mori* chromosomes by Oligopaint fluorescence *in situ* hybridization (FISH). Findings demonstrated that both male Z chromosomes are similar in size and shape. They were also shown to be more compact than either autosomes or the female Z chromosome [13]. Although their results suggest that both male Z chromosomes are repressed, this model is not directly confirmed by comparing the expression levels of genes on different Z chromosomes in males.

In this study, we analyzed allele-specific expression (ASE) by transcriptome analysis using the hybrids (F_1_) between two different *B. mori* strains. Results obtained showed the absence of allelic imbalance in *B. mori* dosage compensation. Furthermore, our results indicate that *B. mori* possesses a nematode-like dosage compensation system, which equally represses both Z chromosomes transcriptionally.

## Materials and Methods

### Silkworm strains

*B. mori* strains, p50T and N4, were maintained in our laboratory. An *od* mutant strain, n03, was provided from Kyushu University with support from the National Bioresource Project (http://silkworm.nbrp.jp). All larvae were reared on mulberry leaves or an artificial diet (SilkMate PS (NOSAN)) under 12 L/12 D conditions at 25°C.

### RNA extraction, molecular sexing and RNA sequence (RNA-Seq)

Embryonic RNAi was performed using F_1_ (N4 female × p50T male) embryos [8,14]. Both DNA and RNA were isolated from the injected embryos at 72 hours post-injection with TRIzol (Thermo Fisher Scientific) [15]. Genomic PCR for molecular sexing was performed using KOD FX Neo (TOYOBO) with the primer sets listed in table S1. RNA extracted from three individuals was pooled together for each experimental group. Libraries for RNA-seq were generated using the Illumina TruSeq RNA Prep Kit (Illumina). The libraries were sequenced by Hiseq 2500 sequencing system (Illumina) with 100 bp paired-end reads.

Adapter sequences and low-quality reads were removed with Trim Galore! (https://www.bioinformatics.babraham.ac.uk/projects/trim_galore/). Genomic short reads of p50T and N4 (NCBI accession numbers DRR064025 and DRR059712) were mapped to *B. mori* reference genome [16] using BWA-MEM [17]. Variant calling was performed using GATK [18]. Obtained variant call format files were loaded onto g2gtools (https://g2gtools.readthedocs.io/) to generate paternal and maternal reference genome sequences. Also, the read counts for each gene corresponding to the paternal and maternal alleles were estimated using EMASE [19] following the authors’ instructions. Additionally, differential expression (DE) analysis was performed using edgeR [20] and the read counts obtained from EMASE. Genes with CPM values less than 1 in both groups were filtered out in DE analysis.

### Observation of larval epidermal cells

A fifth-instar larva was dissected from the ventral side, and its silk glands, tracheae, midgut and fat body were removed. After washing with 1 mL of phosphate-buffered saline (PBS) three times, the integument lining was stained with 100 μL of PBS containing 10 μg/mL Hoechst 33342 (DOJINDO) for 5 min in the dark at room temperature. Also, the muscle fibres were cut, and the sixth dorsal segment was cut out after washing three times with 1 mL of PBS. They were then placed between the slide and coverslip. Larval epidermal cells were observed under fluorescence microscope Axio Zoom.V16 (Zeiss).

## Results

RNA interference (RNAi)-mediated knockdown was performed to investigate the presence of an allelic imbalance *B. mori* dosage compensation. Either *Masc* or control (*GFP*) siRNA was injected into the F_1_ (N4 female × p50T male) embryos (figure 1*a*). Furthermore, RNA-seq analysis was also performed using the total RNA extracted from those embryos. The boxplots of the mapping results showed that the expression levels of Z-linked genes were upregulated in *Masc* knocked down males due to a failure of dosage compensation, whereas such a bias was not observed in females (figure 1*b–c*).

**Figure 1.**
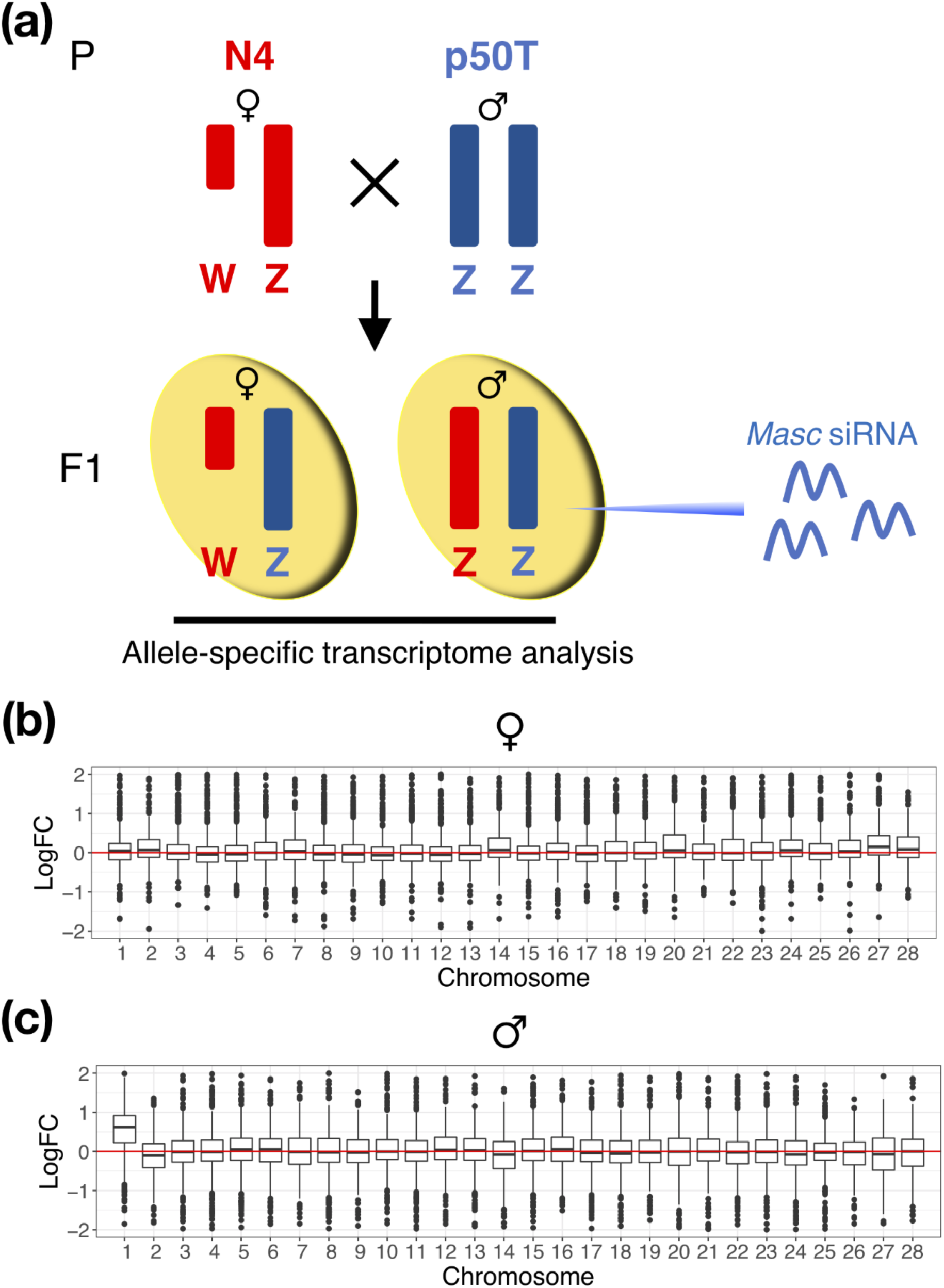
A failure of dosage compensation in *Masc* RNAi embryos. (a) Schematic illustrations of ASE analysis in F_1_ individuals. (b–c) Chromosomal distribution of differentially expressed transcripts in male (b) and female (c) embryos injected with *GFP* (control) or *Masc* siRNAs. logFC: log2-fold change with *Masc* siRNA injected individuals concerning control individuals.

Next, ASE analysis to determine whether each read was transcribed from the maternal or paternal chromosome, followed by a DE analysis (figure 2, table S2, table S3). Our results demonstrated that genes on both maternal and paternal Z chromosomes were upregulated in *Masc* knocked down males but not in females (figure 2*a–c*). Additionally, in *Masc* knocked down males, the upregulated genes were uniformly dispersed throughout the maternal and paternal Z chromosomes (figure 2*a–b*). Most were commonly upregulated in maternal and paternal Z chromosomes (figure S1).

**Figure 2.**
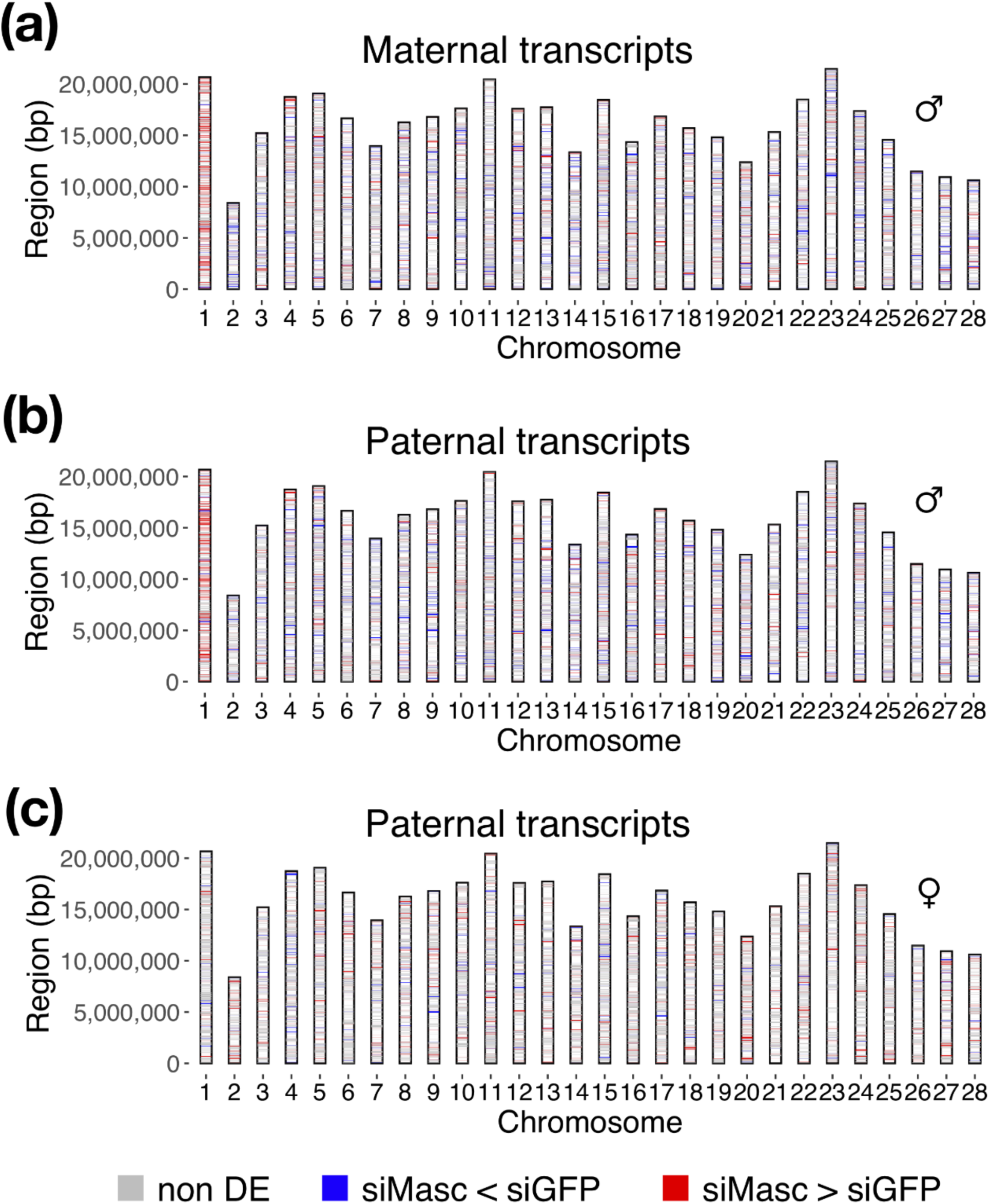
Masc-induced dosage compensation occurs in both maternal and paternal Z chromosomes in males. (a-b) The genome loci of Masc-regulated genes transcribed from maternal (a) and paternal (b) chromosomes in F_1_ males. (c) The genome loci of Masc-regulated genes transcribed from paternal chromosomes in F_1_ females. For each gene on each chromosome, log2-fold change (logFC) with *Masc* siRNA injected individuals (siMasc) concerning control individuals (siGFP) was calculated. The genes whose logFC > 0.5 were coloured in red, logFC < −0.5 were coloured in blue and −0.5 ≦ logFC ≦ 0.5 were coloured in gray. The maternal chromosomes in the F_1_ females were excluded since they possess only a paternal Z chromosome.

Two hypotheses were drawn based on the ASE analysis; they include 1) the transcripts from both male Z chromosomes are balanced as a whole by randomly silencing one of the two Z chromosomes in each cell, and 2) both copies of the male Z chromosome are repressed. To test the hypotheses, uric acid accumulation was monitored in each larval epidermal cell of a male heterozygous by observing Z-linked mutation, *od*, which causes a defect in urate granule formation [21]. The gene responsible for *od* is *B. mori* homolog of mammalian *biogenesis of lysosome-related organelles complex 1, subunit 2* (*BmBLOS2*), which was upregulated in *Masc* siRNA injected males (table S3). This suggests that its expression is regulated by dosage compensation under the control of Masc. The larval epidermal cells of the *od* mutant (*od*/*od*) were translucent due to lack of uric acid accumulation, whereas those of the wild type (*+/+*) were pigmented white (figure S2). Uric acid was uniformly accumulated in all epidermal cells of a male heterozygous for the *od* mutation (*od/+*) like in the wild type (figure S2). The result indicated that *B. mori* males do not randomly silence one of the two Z chromosomes in each cell as eutherians.

## Discussion

Previously, Kiuchi et al. (2014) reported that most Z-chromosome-derived transcripts expressed higher in *Masc* knocked down male embryos are dispersed throughout the Z chromosome [8]. Notably, our results verified that the upregulated genes are dispersed throughout both Z chromosomes in *Masc* knocked down males. It also showed that most of the dispersed genes are commonly upregulated in both Z chromosomes (figure 1*a–b*, figure S1). However, approximately 40.0% of the Z-linked genes were not upregulated (logFC < 0.5) in *Masc* knocked down males (table S3). This group of genes appears to be unaffected by dosage compensation. This variation may be due to the tendency to accumulate male-biased genes on the Z chromosome in species with a WZ sex determination system [22]. Previous studies in several lepidopteran species demonstrated enrichment of male-biased genes on the Z chromosome. Also, a reduction in female-biased genes on Z chromosomes was observed in the abdomen or gonad [9,12,23]. Thus, considerable Masc-dependent male-biased genes on the Z chromosome seem unaffected by dosage compensation.

In this study, ASE analysis was performed in F_1_ embryos, and the results revealed that the genes on both maternal and paternal Z chromosomes were upregulated in *Masc* knocked down males (figure 2*a–c*). Uric acid accumulation was monitored in each larval epidermal cell of an *od/+* male to exclude the possibility that one of the two Z chromosomes was randomly inactivated in each cell. The gene responsible for *od, BmBLOS2*, encodes a subunit involved in urate granule formation in the cytoplasm; it demonstrates an autonomous cell function [21,24]. Therefore, uric acid was uniformly accumulated in the epidermal cells of an *od/+* male as in the wild type (figure S2). This phenotype is different from coat colour mutations in *M. musculus* [2], whose dosage compensation is archived by randomly silencing one of the two X chromosomes in each cell. Therefore, our results showed that *B. mori* establishes nematode-like dosage compensation, which equally suppresses both copies of the male Z chromosome. These results corroborate a recent study showing that both male Z chromosomes are similar in size and shape and more compact than either autosomes or the female Z chromosome using Oligopaint FISH [13].

Intriguingly, *B. mori* and *C. elegans* share a holocentric chromosomal structure. Additionally, Gu et al. (2019) reported that in Monarch butterfly, *Danaus plexippus*, depletion of histone H4 acetylation on lysine 16 (H4K16ac) on the ancestral Z segment (which is also the Z in other lepidopteran insects) is associated with dosage compensation [25]. Although there are some differences, depletion of H4K16ac on the X chromosome is also associated with dosage compensation in *C. elegans* [26]. These observations show the similarities between Lepidoptera and *C. elegans*, notwithstanding that they belong to distinct clades and have different sex determination systems. In *C. elegans*, a member of condensin complexes, condensin I^DC^, binds to X chromosomes and contributes to dosage compensation [27–30]. Some condensin subunits have been repeatedly and independently lost in multiple insect orders, including Lepidoptera. Their loss has no relationship with pairing prevalence [31]. These observations suggest that the condensin complexes may have acquired new functions in Lepidoptera. Hence, further studies on the holocentric chromosome structure [32] and the functions of condensin complexes in *B. mori* are recommended to reveal the detailed mechanisms of dosage compensation in Lepidoptera.

## Acknowledgements

We thank the Institute for Sustainable Agro-ecosystem Services, The University of Tokyo, for facilitating the mulberry cultivation and the Biotron Facility at the University of Tokyo for rearing the silkworms. We thank Drs. Hiroyuki Hikida, Keisuke Shoji, and Toru Shimada for their useful comments. We are grateful to Wakako Saito and Natsuki Nakashima for technical assistance for silkworm maintenance. This work was supported by JSPS/MEXT KAKENHI grant numbers JP15H02482 and JP17H06431 to SK and TK.

## Author contributions

TK and SK designed the study. TK prepared insects used in this study. KT and TK performed molecular experiments. KT and MK analyzed the data. YS prepared the RNA-seq library and performed Hiseq 2500 sequencing experiments. KT wrote the draft manuscript and TK revised it with the inputs of SK.

## Data accessibility

RNA-seq data was submitted to DDBJ under the accession number DRA013569.

## Figures and legends

**Figure S1.**
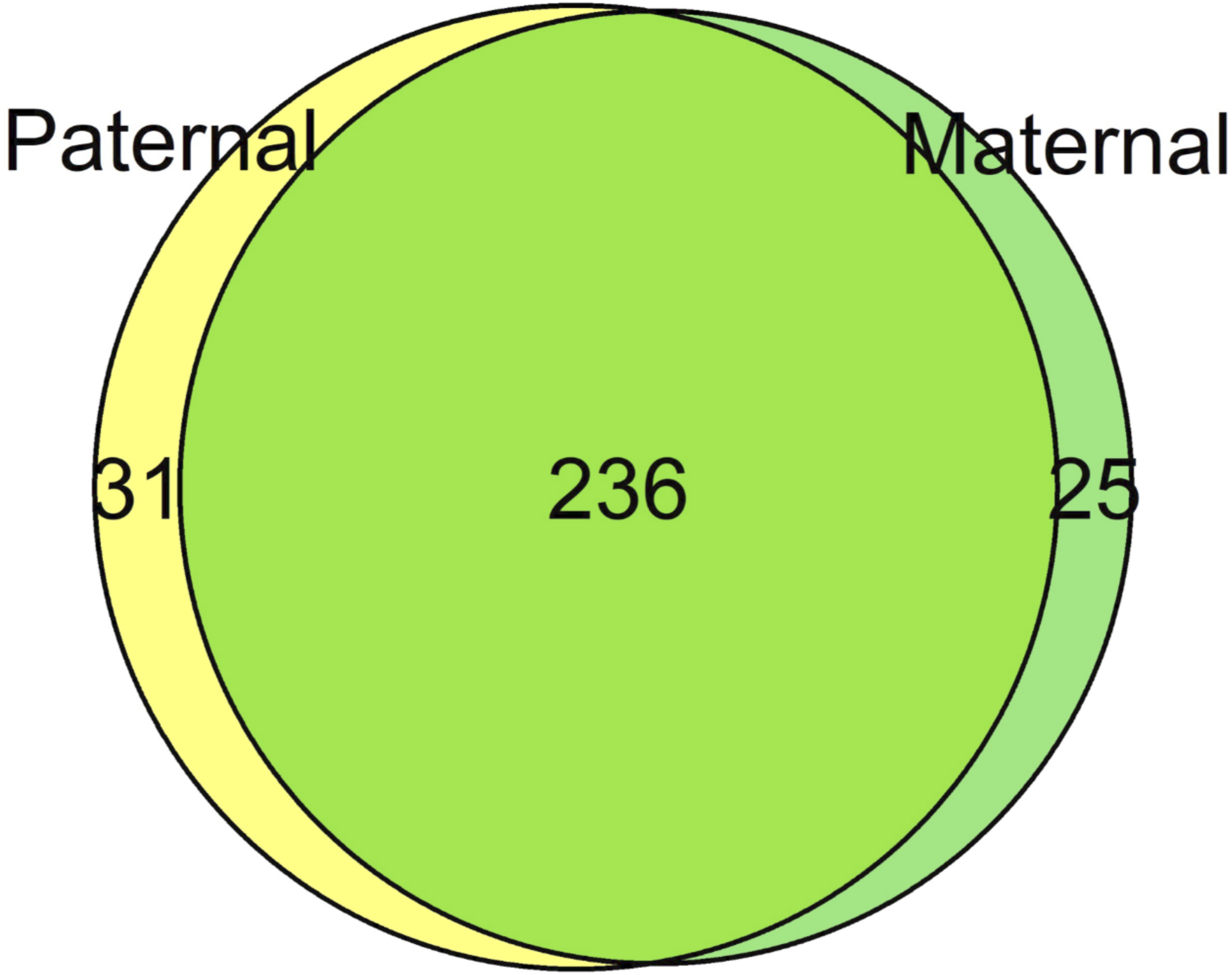
Commonly upregulated genes in maternal and paternal Z chromosomes in F_1_ males by *Masc* mRNA depletion. The green and yellow circles indicate Z-linked genes whose expression was upregulated (logFC > 0.5) in *Masc* siRNA injected males in the maternal and paternal alleles, respectively.

**Figure S2.**
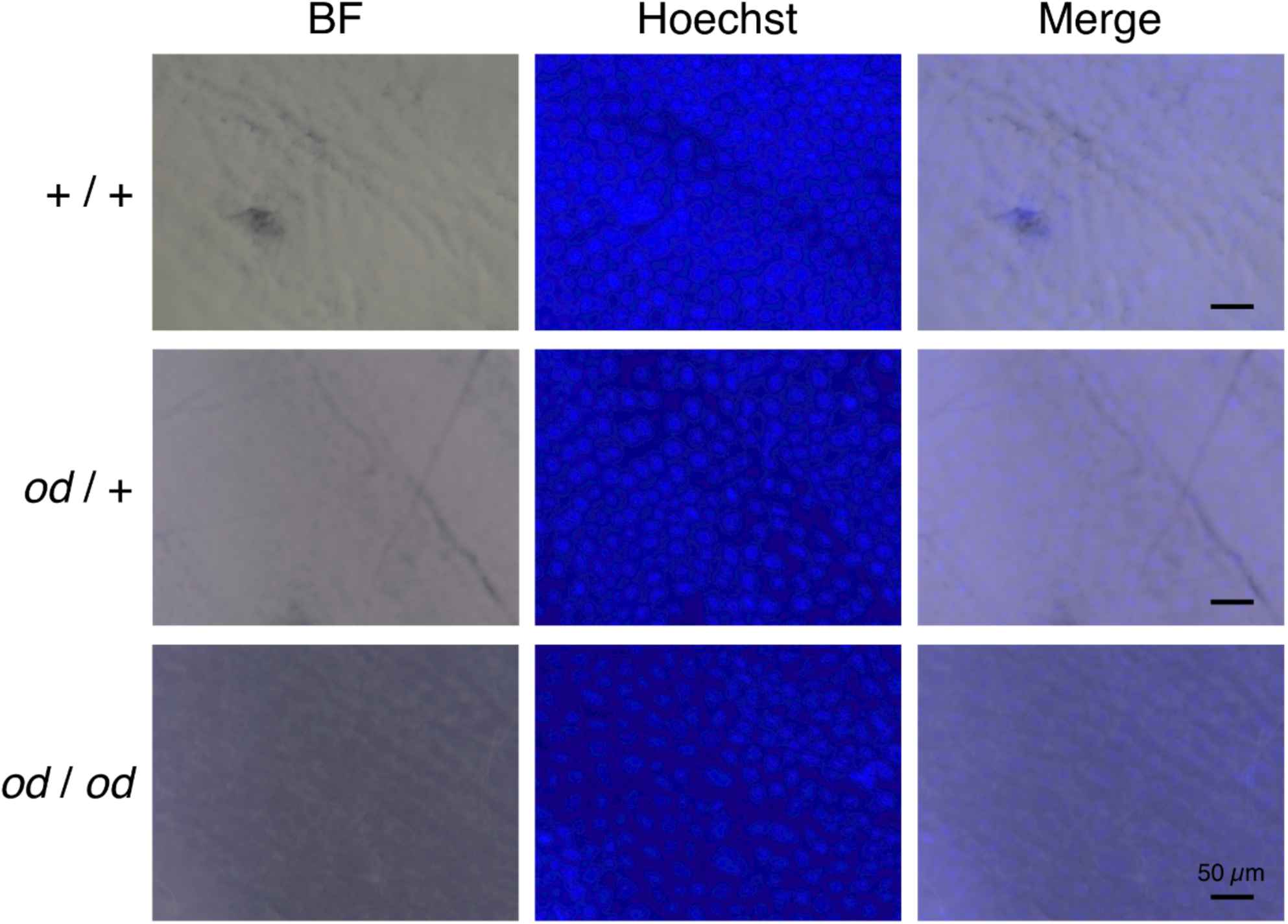
Uric acid accumulation of larval epidermal cells in the *od*/*+* male showed both male Z chromosomes were repressed. Microscopic observation of larval epidermal cells in *+*/*+* (top), *od*/*+* (middle) and *od*/*od* (bottom) males. Bright-field (BF; left); Hoechst staining (center); merge (right). Scale bars, 50 μm.

**Table S1**. Primer and RNA sequences used in this study.

**Table S2**. The result of ASE analysis in F_1_ females. DE analysis between *Masc* siRNA injected and control embryos was performed for each parental allele. logCPM: average log count per million, FDR: false discovery rate, TPM: transcripts per million.

**Table S3**. The result of ASE analysis in F_1_ males. DE analysis between *Masc* siRNA injected and control embryos was performed for each parental allele. The abbreviation was described in table S2.

## References

1. Lyon MF. 1961 Gene Action in the X-chromosome of the Mouse (Mus musculus L.). Nature 190, 372–373. (doi:10.1038/190372a0)

2. Frasee AS, Sobey S, Spicer CC. 1953 Mottled, a sex-modified lethal in the house mouse. J. Genet. 51, 217–221. (doi:10.1007/BF03023293)

3. Whitworth DJ, Pask AJ. 2016 The X factor: X chromosome dosage compensation in the evolutionarily divergent monotremes and marsupials. Semin. Cell Dev. Biol. 56, 117–121. (doi:10.1016/J.SEMCDB.2016.01.006)

4. Hamada FN, Park PJ, Gordadze PR, Kuroda MI. 2005 Global regulation of X chromosomal genes by the MSL complex in Drosophila melanogaster. Genes Dev. 19, 2289–2294. (doi:10.1101/GAD.1343705)

5. Straub T, Gilfillan GD, Maier VK, Becker PB. 2005 The Drosophila MSL complex activates the transcription of target genes. Genes Dev. 19, 2284–2288. (doi:10.1101/GAD.1343105)

6. Ercan S, Lieb JD. 2009 C. elegans dosage compensation: A window into mechanisms of domain-scale gene regulation. Chromosom. Res. 17, 215–227. (doi:10.1007/S10577-008-9011-0/FIGURES/3)

7. Tanaka Y. 1916 Genetic studies on the silkworm. J. Coll. Agric. Tohoku Imp. Univ. Sapporo, Japan 7, 129–255.

8. Kiuchi T et al. 2014 A single female-specific piRNA is the primary determiner of sex in the silkworm. Nature 509, 633–636. (doi:10.1038/nature13315)

9. Gu L, Walters JR, Knipple DC. 2017 Conserved patterns of sex chromosome dosage compensation in the Lepidoptera (WZ/ZZ): Insights from a moth neo-Z chromosome. Genome Biol. Evol. 9, 802–816. (doi:10.1093/gbe/evx039)

10. Huylmans AK, Macon A, Vicoso B. 2017 Global dosage compensation os ubiquitous in Lepidoptera, but counteracted by the masculinization of the Z chromosome. Mol. Biol. Evol. 34, 2637–2649. (doi:10.1093/molbev/msx190)

11. Smith G, Chen Y-R, Blissard GW, Briscoe AD. 2014 Complete dosage compensation and sex-biased gene expression in the moth Manduca sexta. Genome Biol. Evol. 6, 526–537. (doi:10.1093/gbe/evu035)

12. Walters JR, Hardcastle TJ, Jiggins CD. 2015 Sex chromosome dosage compensation in Heliconius butterflies: Global yet still incomplete? Genome Biol. Evol. 7, 2545–2559. (doi:10.1093/gbe/evv156)

13. Rosin LF, Chen Y, Lei EP. 2021 Dosage compensation in Bombyx mori is achieved by partial repression of both Z chromosomes in males. bioRxiv, 2021.07.18.452821. (doi:10.1101/2021.07.18.452821)

14. Kiuchi T, Katsuma S. 2022 Functional Characterization of Silkworm PIWI Proteins by Embryonic RNAi. Methods Mol. Biol. 2360, 19–31. (doi:10.1007/978-1-0716-1633-8_3)

15. Kiuchi T, Katsuma S. 2022 Functional Characterization of Silkworm PIWI Proteins by Embryonic RNAi. Methods Mol. Biol. 2360, 19–31. (doi:10.1007/978-1-0716-1633-8_3)

16. Kawamoto M et al. 2019 High-quality genome assembly of the silkworm, Bombyx mori. Insect Biochem. Mol. Biol. 107, 53–62. (doi:10.1016/j.ibmb.2019.02.002)

17. Li H. 2013 Aligning sequence reads, clone sequences and assembly contigs with BWA-MEM., 1303.3997 [q-bio.GN].

18. McKenna A et al. 2010 The genome analysis toolkit: A MapReduce framework for analyzing next-generation DNA sequencing data. Genome Res. 20, 1297–1303. (doi:10.1101/gr.107524.110)

19. Raghupathy N, Choi K, Vincent MJ, Beane GL, Sheppard KS, Munger SC, Korstanje R, Pardo-Manual de Villena F, Churchill GA. 2018 Hierarchical analysis of RNA-seq reads improves the accuracy of allele-specific expression. Bioinformatics 34, 2177–2184. (doi:10.1093/bioinformatics/bty078)

20. Robinson MD, McCarthy DJ, Smyth GK. 2010 edgeR: a Bioconductor package for differential expression analysis of digital gene expression data. Bioinformatics 26, 139–140. (doi:10.1093/bioinformatics/btp616)

21. Fujii T, Daimon T, Uchino K, Banno Y, Katsuma S, Sezutsu H, Tamura T, Shimada T. 2010 Transgenic analysis of the BmBLOS2 gene that governs the translucency of the larval integument of the silkworm, Bombyx mori. Insect Mol. Biol. 19, 659–667. (doi:10.1111/j.1365-2583.2010.01020.x)

22. Grath S, Parsch J. 2016 Sex-biased gene expression. Annu. Rev. Genet. 50, 29–44. (doi:10.1146/ANNUREV-GENET-120215-035429)

23. Arunkumar KP, Mita K, Nagaraju J. 2009 The silkworm Z chromosome is enriched in testis-specific genes. Genetics 182, 493–501. (doi:10.1534/genetics.108.099994)

24. Takasu Y, Kobayashi I, Beumer K, Uchino K, Sezutsu H, Sajwan S, Carroll D, Tamura T, Zurovec M. 2010 Targeted mutagenesis in the silkworm Bombyx mori using zinc finger nuclease mRNA injection. Insect Biochem. Mol. Biol. 40, 759–765. (doi:10.1016/J.IBMB.2010.07.012)

25. Gu L, Reilly PF, Lewis JJ, Reed RD, Andolfatto P, Walters JR. 2019 Dichotomy of dosage compensation along the neo Z chromosome of the monarch butterfly. Curr. Biol. 29, 4071-4077.e3. (doi:10.1016/j.cub.2019.09.056)

26. Wells MB, Snyder MJ, Custer LM, Csankovszki G. 2012 Caenorhabditis elegans dosage compensation regulates histone H4 chromatin state on X chromosomes. Mol. Cell. Biol. 32, 1710–1719. (doi:10.1128/MCB.06546-11/ASSET/A8133109-DBEE-4F2C-BE2E-B7C4F325538A/ASSETS/GRAPHIC/ZMB9991094730007.JPEG)

27. Lieb JD, Albrecht MR, Chuang P-T, Meyer BJ. 1998 MIX-1: An essential component of the C. elegans mitotic machinery executes X chromosome dosage compensation. Cell 92, 265–277. (doi:10.1016/S0092-8674(00)80920-4)

28. Chuang PT, Albertson DG, Meyer BJ. 1994 DPY-27: A chromosome condensation protein homolog that regulates C. elegans dosage compensation through association with the X chromosome. Cell 79, 459–474. (doi:10.1016/0092-8674(94)90255-0)

29. Tsai CJ, Mets DG, Albrecht MR, Nix P, Chan A, Meyer BJ. 2008 Meiotic crossover number and distribution are regulated by a dosage compensation protein that resembles a condensin subunit. Genes Dev. 22, 194–211. (doi:10.1101/GAD.1618508)

30. Lieb JD, Capowski EE, Meneely P, Meyer BJ. 1996 DPY-26, a link between dosage compensation and meiotic chromosome segregation in the nematode. Science 274, 1732–1736. (doi:10.1126/SCIENCE.274.5293.1732)

31. King TD, Leonard CJ, Cooper JC, Nguyen S, Joyce EF, Phadnis N. 2019 Recurrent losses and rapid evolution of the condensin II complex in insects. Mol. Biol. Evol. 36, 2195–2204. (doi:10.1093/MOLBEV/MSZ140)

32. Drinnenberg IA, deYoung D, Henikoff S, Malik HS ing. 2014 Recurrent loss of CenH3 is associated with independent transitions to holocentricity in insects. Elife 3. (doi:10.7554/ELIFE.03676)

